# Predicting age and clinical risk from the neonatal connectome

**DOI:** 10.1101/2020.09.28.317180

**Authors:** Yassine Taoudi-Benchekroun, Daan Christiaens, Irina Grigorescu, Oliver Gale-Grant, Andreas Schuh, Maximilian Pietsch, Andrew Chew, Nicholas Harper, Shona Falconer, Tanya Poppe, Emer Hughes, Jana Hutter, Anthony N Price, J-Donald Tournier, Lucilio Cordero-Grande, Serena J Counsell, Daniel Rueckert, Tomoki Arichi, Joseph V Hajnal, A David Edwards, Maria Deprez, Dafnis Batalle

**Affiliations:** Centre for the Developing Brain, School of Imaging Sciences & Biomedical Engineering, King’s College London, London, United Kingdom; Department of Electrical Engineering, ESAT/PSI, KU Leuven, Leuven, Belgium; Department of Forensic and Neurodevelopmental Science, Institute of Psychiatry, Psychology and Neuroscience, King’s College London, London, United Kingdom; Biomedical Image Analysis Group, Department of Computing, Imperial College London, London, United Kingdom; Biomedical Image Technologies, ETSI Telecomunicación, Universidad Politécnica de Madrid & CIBER-BBN, Madrid, Spain; Institute for Artificial Intelligence and Data Science in Medicine and Healthcare, Faculty of Medicine and Informatics, Technical University of Munich, Munich, Germany; Department of Bioengineering, Imperial College London, London, United Kingdom; Children’s Neurosciences, Evelina London Children’s Hospital, Guy’s and St Thomas’ NHS Trust, London, United Kingdom; MRC Centre for Neurodevelopmental Disorders, King’s College London, London, United Kingdom

## Abstract

The development of perinatal brain connectivity underpins motor, cognitive and behavioural abilities in later life. Diffusion MRI allows the characterisation of subtle inter-individual differences in structural brain connectivity. Individual brain connectivity maps (connectomes) are by nature high in dimensionality and are complex to interpret. Machine learning methods are a powerful tool to uncover properties of the connectome which are not readily visible, and can give us clues as to how and why individual developmental trajectories differ.

In this manuscript we used Deep Neural Networks and Random Forests to predict demographic and neurodevelopmental characteristics from neonatal structural connectomes in a large sample of babies (n = 524) from the developing Human Connectome Project. We achieved an accurate prediction of post menstrual age (PMA) at scan in term-born infants (Mean absolute error (MAE) = 0.72 weeks, r = 0.83 and p<0.001). We also achieved good accuracy when predicting gestational age at birth in a cohort of term and preterm babies scanned at term equivalent age (MAE = 2.21 weeks, r = 0.82, p<0.001). We subsequently used sensitivity analysis to obtain feature relevance from our prediction models, with the most important connections for prediction of PMA and GA found to be predominantly thalamocortical. From our models of PMA at scan for infants born at term, we computed a brain maturation index (*predicted age minus actual age*) of individual preterm neonates and found a significant correlation between this index and motor outcome at 18 months corrected age. Our results demonstrate the applicability of machine learning techniques in analyses of the neonatal connectome, and suggest that a neural substrate for later developmental outcome is detectable at term equivalent age.

## 1. INTRODUCTION

Magnetic Resonance Imaging (MRI) allows a broad range of in vivo insights about the structure and function of the human brain. Diffusion MRI in particular enables the characterization of microstructural changes in the orientation and organisation of major white matter tracts, the systematic description of whole-brain structural networks: the human *connectome* (Honey et al., 2010; Sporns et al., 2005).

During the perinatal period, the brain undergoes significant changes and consolidation of structural connectivity, which are thought to underpin the expansion of motor, cognitive and behavioural abilities (Johnson, 2001). Since the inception of *connectomics* (Hagmann 2005; Sporns, Tononi, and Kötter 2005) several studies have tried to characterise early development of the structural connectome (Fan et al. 2011; Hagmann et al. 2010). Subtle alterations in the development of brain connectivity have been suggested to underlie atypical neurodevelopmental outcome in populations with perinatal risk factors, such as children born preterm (Batalle et al., 2018). Preterm birth comprises approximately 11% of all births, and is the main global cause of death and disability in children under 5 years of age (Blencowe et al., 2012), as well as representing one of the most pervasive perinatal risk factors for atypical neurodevelopment (Wood et al., 2000). It has been associated with an increased risk of developing neurodevelopmental conditions such as motor, visuospatial and sensorimotor delay (Marlow et al., 2007), inattention, anxiety and social difficulties (Johnson and Marlow, 2014), autism spectrum (Johnson et al., 2010), cerebral palsy (Marlow et al., 2005) or psychiatric disorders in adulthood such as depression and bipolarity (Nosarti et al., 2012). Therefore, improving our understanding of how preterm birth affects structural brain development remains an important goal.

With this goal in mind, the developing Human Connectome Project (dHCP) has collected demographic and MRI data from a large cohort of term and preterm-born neonates. The dHCP dataset comprises structural, diffusion and functional MRI data with high spatial, angular, and temporal resolutions. Features of this project include: advances in hardware (Hughes et al., 2017) and protocols for neonatal diffusion MRI acquisition (Hutter et al. 2018); the use of multiband techniques to accelerate acquisition time combined with approaches to correct motion (Cordero-Grande et al. 2016; Cordero-Grande et al. 2019; Christiaens et al. 2021); and the development of state-of-the-art preprocessing pipelines for neonatal MRI data (Bozek et al., 2018; Christiaens et al. 2018; Bastiani et al. 2019; Fitzgibbon et al. 2020; Makropoulos et al., 2018). Together, these have significantly improved neonatal MRI acquisition and reconstruction methods.

Despite the resultant advances in data quality, studying the neonatal connectome remains challenging. Indeed, many methodological issues hamper the interpretation of the connectome (Sporns, 2013) including the difficulty of detecting origins and termination of connections (Jbabdi and Johansen-Berg, 2011) and returning a high number of false positive streamlines (Maier-Hein et al., 2017). However, the development of state-of-the-art diffusion MRI pipelines have partially addressed some of these issues (Makropoulos et al., 2018; Christiaens et al., 2021), and machine learning approaches are now well suited to discover complex underlying patterns in the structural connectome. Deep neural networks, which are known for their ability to model complex non-linear multivariate relationships, can help uncover hidden patterns in the connectome, and potentially detect atypical patterns of connectivity in individual subjects.

In adult brain connectivity research, a number of studies have used machine learning and deep learning to study the structural connectome (see (Brown and Hamarneh, 2016) for a summary). Some studies also used similar approaches in neonates: Kawahara and colleagues have developed BrainNetCNN, a Convolutional Neural Network composed of edge to edge, edge to node, and node to graph convolution filters on structural connectivity to predict post menstrual age (PMA) at scan and cognitive performance from the structural connectome (Kawahara et al., 2017). A recent study by Girault and colleagues similarly focused on using the structural connectome at birth to predict cognitive abilities (Mullen score) at age 2 with dense neural networks (Girault et al., 2019). However, little is known about the predictive power of the connectome in a large normative neonatal population such as that of the dHCP.

A promising method in adult and neonatal neuroscience is the study of the “brain maturation index” (also known as “brain delta” or “predicted age difference”) corresponding to the apparent age of the subject as compared to the norm (Dosenbach et al. 2010; Cao et al. 2015; Jonsson et al. 2019; Liem et al. 2017; Smith et al. 2019). By training regression models to fit the age of subjects from large normative imaging datasets, we can predict the age of individual subjects and compute the difference between the prediction and subject’s true age. This difference gives information about brain maturation and its divergence from the population norm. As such, in adults, a positive (predicted age > true age) difference is interpreted as demonstrating accelerated ageing, and is associated with disorders such as cognitive impairment (Liem et al., 2017), schizophrenia (Koutsouleris et al., 2014) or diabetes (Franke et al., 2013). In neurodevelopment, studying the brain maturation index is therefore extremely relevant for preterm born infants where neurodevelopmental delays and psychiatric disorders often occur (Brown et al., 2017; Galdi et al., 2020; Rasmussen et al., 2017).

In this work, we propose the use of two different machine learning algorithms - Random Forests (RF) and Deep Neural Networks (DNN) - to predict PMA at scan (i.e. brain maturation), and gestational age (GA) at birth (i.e. degree of prematurity), from the neonatal structural connectome in a large sample of neonates scanned at term equivalent age. Additionally, we use sensitivity analysis to obtain feature relevance in the prediction models thereby identifying the connections and brain regions impacting the predictions. Finally, using models obtained for the term-born cohort, brain maturation index was computed for our preterm-born cohort and subsequently correlated to neurodevelopment at 18 months.

## 2. Methods and Materials

### 2.1 Participants

All participants were part of the dHCP, approved by the National Research Ethics Service West London committee (14/LO/1169).

524 infants (240 female and 284 male), born between 23^+0^ weeks and 42^+2^ week of gestation, underwent MRI between 37^+1^ weeks and 45^+1^ weeks as part of the dHCP (2^nd^ data release). The participant gestational age at birth (GA) and postmenstrual age at scan (PMA) distribution is presented in Figure 1A-B. Full participant clinical information is presented in Table 1.

**Table 1.** Detailed sample and outcome characteristics.

|  | Term Born<br>N = 418 | Preterm born<br>N = 106 | p-value* |
| --- | --- | --- | --- |
| Gestational age at birth [weeks<sup>+days</sup>] | Median = 40<sup>+1</sup><br>IQR = 39<sup>+0</sup> – 40<sup>+6</sup> | Median = 32<sup>+2</sup><br>IQR = 28<sup>+5</sup> – 34<sup>+4</sup> | <0.001 |
| Postmenstrual age at scan [weeks<sup>+days</sup>] | Median = 41<sup>+0</sup><br>IQR = 39<sup>+6</sup> – 42<sup>+2</sup> | Median = 41<sup>+0</sup><br>IQR = 39<sup>+5</sup> – 42<sup>+2</sup> | 0.771 |
| Sex, no. of female (%) | 195 (47%) | 45 (42%) | 0.439 |
| BSID-III, no. (% of total population) | 322 (77%) | 77 (73%) | 0.343 |
| Gestational age at birth (weeks<sup>+days</sup>) of subjects with BSID-III data available | Median = 39<sup>+6</sup><br>IQR = 39<sup>+1</sup> – 40<sup>+5</sup> | Median = 31<sup>+6</sup><br>IQR = 28<sup>+4</sup> – 34<sup>+5</sup> | < 0.001 |
| Postmenstrual age at scan (weeks<sup>+days</sup>) of subjects with BSID-III data available | Median = 41<sup>+1</sup><br>IQR = 39<sup>+6</sup> – 42<sup>+2</sup> | Median = 41<sup>+1</sup><br>IQR = 39<sup>+5</sup> – 42<sup>+3</sup> | 0.957 |
| IMD Score of subjects with BSID-III data available | Mean = 13839<br>STD = 7387 | Mean = 16465<br>STD = 7805 | 0.005 |
| Fine Motor | Mean = 20.69<br>STD = 3.12 | Mean = 20.22<br>STD = 3.36 | 0.241 |
| Gross Motor | Mean = 17.34<br>STD = 2.37 | Mean = 16.75<br>STD = 2.40 | 0.052 |
| Cognitive Score | Mean = 10.08<br>STD = 2.15 | Mean = 9.79<br>STD = 2.59 | 0.303 |
| Expressive communication | Mean = 17.26<br>STD = 3.62 | Mean = 16.89<br>STD = 4.03 | 0.434 |
| Receptive Communication | Mean = 18.36<br>STD = 4.29 | Mean = 18.51<br>STD = 5.26 | 0.800 |

\*p-values computed with two tailed independent t-test or chi-square test as appropriately.

**Figure 1.**
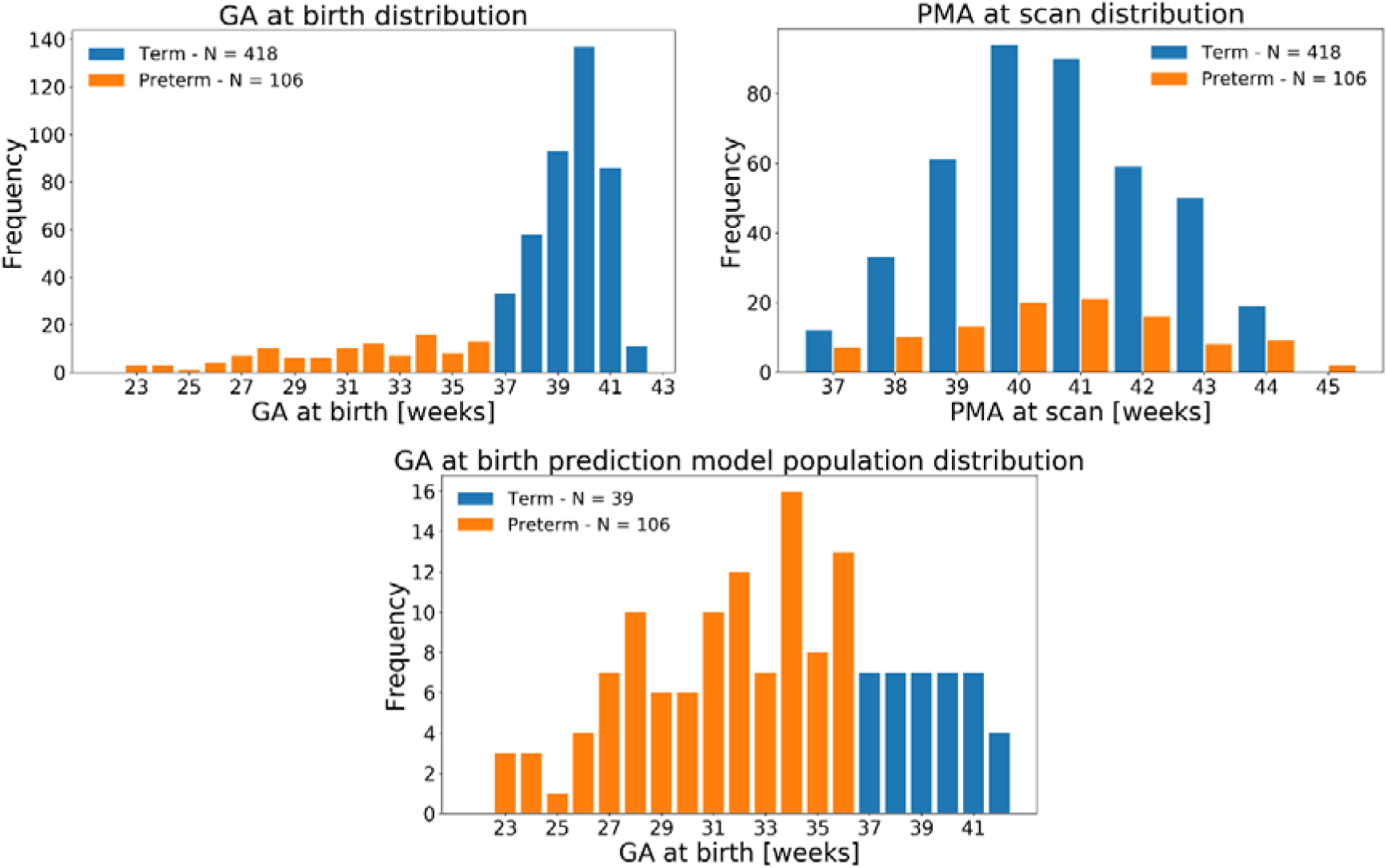
Distribution of (A) GA at birth and (B) PMA at scan of full cohort (N=524). (C) GA at birth of cohort used for predicting GA at birth.

The Bayley III Scales of Infant and Toddler Development (BSID-III) (Bayley 2006) were collected at 18 months corrected age and available for 314 infants including 50 preterm-born infants. We used scores of motor (fine and gross), communication (expressive and receptive) and cognitive (raw) score. Assessments were carried out by experienced paediatricians or psychologists. Detailed assessment distributions are presented in Table 1.

### 2.2 MRI acquisition

All scans were collected in the Evelina Newborn Imaging Centre based on the Neonatal Intensive Care Unit, St Thomas hospital London using a Philips Achieva 3T scanner (Best, NL) and the dHCP neonatal brain imaging system which includes a 32 channel receive neonatal head coil (Rapid Biomedical GmbH, Rimpar, DE) (Hughes et al., 2017). Informed written parental consent was obtained prior to imaging. Positioning of all infants was done with a lightweight protective “shell”, which was positioned on an MRI safe trolley to ease transportation. Immobilization of the infants in the shell was done using bead filled inflatable pads (Pearltec, Zurich, CH). In addition to the pads, acoustic protection included earplugs moulded from a silicone-based putty (President Putty, Coltene Whaledent, Mahwah, NJ, USA) placed in the external auditory meatus and neonatal earmuffs (MiniMuffs, Natus Medical Inc, San Carlos, CA, USA). To avoid sudden sound changes which might wake up the infant, the MRI software was modified in order to gradually increase the noise from 0 to the average operating point (Hughes et al., 2017). All scans were supervised by a paediatrician or neonatal nurse experienced in MRI procedures; vital signs including pulse oximetry, temperature and electrocardiography data were monitored throughout data acquisition. All infants were scanned during natural unsedated sleep following feeding.

T2-weighted images were acquired using a Turbo spin echo sequence with parameters TR = 12s and TE = 156ms, SENSE factor 2.11 (axial) and 2.54 (sagittal) with overlapping slices (resolution = 0.8 × 0.8 × 1.6 mm^3^). Super-resolution methods (Kuklisova-Murgasova et al., 2012) as well as motion correction methods (Cordero-Grande et al., 2018) were combined to maximise precision and resolution of T2-weighted images (resolved to 0.8 × 0.8 × 0.8 mm^3^). Diffusion weighted imaging was acquired in 300 directions with parameters TR = 3.8s, TE = 90ms, SENSE factor 1.2, multiband factor 4, resolution = 1.5 × 1.5 × 3mm^3^ (with 1.5mm slice overlap), diffusion gradient encoding: b=0 s/mm (n=20), b=400 s/mm (n=64), b=1000 s/mm (n=88), b=2600 s/mm (n=128), and using interleaved phase encoding (Hutter et al., 2018b).

### 2.3 Pre-processing and connectome generation

Tissue segmentation of T2-weighted volumes was performed using a neonatal specific segmentation pipeline (Makropoulos et al., 2014) and template (Schuh et al. 2018). Parcellation of 90 cortical and subcortical regions (Shi et al., 2011) adapted to the dHCP weekly age-dependant high-resolution bespoke template (Schuh et al. 2018) was propagated to each subject’s T2w native space through non-linear registration based in a diffeomorphic symmetric image normalization method (SyN) available in ANTS software (Avants et al., 2011), using T2w contrast and tissue segmentation as input channels. Tissue maps and atlas parcellation were propagated from each T2w native space to each subject’s diffusion native space with a rigid registration using b=0 volumes as target. All rigid registrations were performed with IRTK software (Schnabel et al. 2001). Details of the 90 cortical and subcortical regions are presented in Supplementary Table 1.

Diffusion MRI was reconstructed at an effective resolution of 1.5mm isotropic and denoised using a patch-based estimation of the diffusion signal based on random matrix theory (Veraart et al., 2016). Gibbs ringing was suppressed (Kellner et al., 2016) and B0 field map estimated from b=0 volumes in order to correct magnetic susceptibility-induced distortion using FSL Topup (Andersson et al., 2003). Data was corrected for slice-level motion and distortion in a data-driven q-space representation using a bespoke spherical harmonics and radial decomposition (SHARD) basis of rank 89 corresponding to spherical harmonics of order lmax=0,4,6,8 for each respective shell, with registration operating at a reduced rank of 22 (Christiaens et al., 2018). DWI intensity inhomogeneity field correction was performed using the ANTs N4 algorithm (Tustison et al., 2010). Tools and pipelines implemented in MRtrix3 (Tournier et al., 2019) were used for quantitative analysis of the diffusion MRI data. Developing neonatal brain tissue undergoes rapid changes in cellular properties and water content that can be to a first approximation captured by a non-negative linear combination of anisotropic signal from relatively mature WM and from isotropic free fluid (Pietsch et al., 2019). We use data from 20 healthy full term control babies from our sample to extract a set of two representative WM (Tournier et al., 2013) and fluid-like (Dhollander et al., 2016, Dhollander et al., 2018) signal fingerprint (*response functions*) that are used to deconvolve each subject’s diffusion signal into a fibre orientation distribution (FOD) image, capturing WM-GM-like signal, and scalar fluid density image using the multi-tissue multi-shell constrained spherical deconvolution technique (Jeurissen et al. 2014). Residual intensity inhomogeneity was corrected, and component densities calibrated using a multi-tissue log-domain intensity normalisation (Raffelt, et al. 2017). Resulting normalised WM-GM-like FODs were used to generate 10 million streamlines with an anatomically constrained probabilistic tractography (ACT) (Smith et al., 2012) with biologically accurate weights (SIFT2) (Smith, et al. 2015). The fibre density SIFT2 proportionality coefficient (μ) for each subject was obtained to achieve inter-subject connection density normalisation. The structural connectivity network of each infant was then constructed by calculating the μ × SIFT2-weighted sum of streamlines connecting each pair of regions (thus built as a symmetric adjacency matrix of size 90×90).

In addition, we used 73 structural connectivity matrices obtained from an independent dataset (Batalle et al., 2017) to test the design of the initial hyperparameters and architecture for predictive algorithms presented in sections 2.4.3 and 2.4.4.

### 2.4 Prediction of age at scan and age at birth

All analyses on this section were performed using Python 3.7. The machine learning library Scikit Learn (Pedregosa et al., 2011) was used for training the RF algorithm. The deep learning framework Keras (version 2.0.3) (Chollet et al., 2015) was used to train the deep learning models.

#### 2.4.1 Feature set

As the structural connectome is presented as a symmetric adjacency matrix (in our case of size 90×90, with 90 brain regions) the lower triangle of the matrix contains all information. We thus extracted and reshaped the lower triangle of each subject’s structural connectome *S*_*i*_ as a 1D vector *X*_*i*_ with number of connectivity elements n = 4005, thus leading to the ensemble *X* of connectivity vectors across *N* subjects:

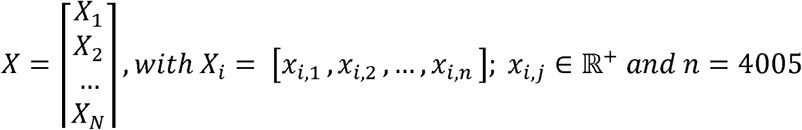

We normalized each data point across the training sets, and normalized the testing set with the training normalization values using a min-max normalization:

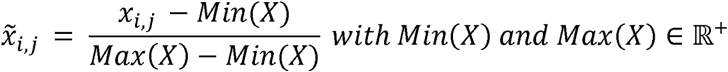

Thus, assuming that testing data also falls between previous ranges, our training and testing data has the following form:

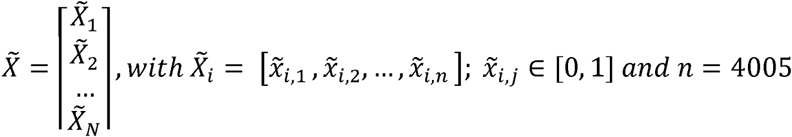

#### 2.4.2 Prediction models

We carried a set of predictions of demographic information on different population samples using different regression algorithms. In each case, we fitted a regression model *f* to predict a variable *Y* representing demographic information (e.g. GA at birth or PMA at scan) for subjects in our dataset. We thus had prediction *Y*’ as follows:

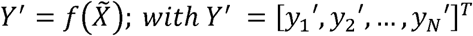

We computed the regressor *f* that minimizes | *Y*′ − *Y* | ^2^. We used two different supervised machine learning regression algorithms to do this: RF and DNN.

#### 2.4.3. Random Forests regression

RF are an ensemble learning method for classification and regression based on constructing a multitude of decision trees (*weak learners*) which are individually trained through the technique of “bagging”. RF makes predictions by averaging the prediction of each individual tree, hence acting as a *strong learner* (Breiman L., 2001). For optimal performance, two main hyperparameters should be tuned: the number of trees (estimators) in the forest and the maximum depth of each tree. The number of trees determines the smoothness of the decision boundary, and the depth corresponds to the maximum number of levels allowed for each tree. RF regressors’ performance often depends on finding the optimal value for these to ensure that there is no overfitting or underfitting.

Here we use the RF regressor from the Scikit Learn *RandomForestRegressor* implementation (Pedregosa et al., 2011). The RF were trained using mean squared error (MSE) as loss function. Hyperparameters were tuned separately for the PMA at scan and GA at birth prediction by performing a grid search on a set of 73 structural connectomes from an independent dataset (Batalle et al., 2017). This allowed us to choose optimal parameters without overfitting our model to the studied data. Hyperparameters used are presented in sections 2.4.6 and 2.4.7.

#### 2.4.4. Deep Neural Networks regression

Deep (Fully Connected) Neural Networks (DNN) are universal function approximators whose parameters can be trained to model complex nonlinear relationships between features and labels via backpropagation (Rumelhart et al., 1986). However, the performance of a DNN also depends on the hyperparameters: design choices are mainly related to the architecture of the network (the layer types, number of layers and number of nodes per layers, activation functions at each layer), the loss function, and the training method (the number of epochs, the optimization function and its parameters).

The DNN in this work were implemented using the deep learning library Keras (version 2.0.3) (Chollet et al., 2015). As performing a grid search to find the best model hyperparameters is computationally expensive when training DNN, we started with a basic architecture built from previous work and common DNN knowledge (Smith 2018), and subsequently optimized these via manual refinement architecture search. To avoid overfitting the model to the data used in this paper, this was done on the same set of 73 structural connectomes from an independent sample as was used for the RF training, independently for the GA at birth and PMA at scan prediction. For both prediction tasks, the models were trained using MSE as a loss function and the Adam optimizer (Kingma and Ba, 2015), albeit with different learning rates. Further details on the network architecture are included in sections 2.4.6 and 2.4.7.

#### 2.4.5 Training and evaluation of the models

To assess the performance of the prediction models, the evaluation metric was calculated on test data excluded from training and hyperparameter tuning. We split the dataset into k groups (folds) and fit the model k times. Each time, one group is used to evaluate performance, while the rest of the groups are used for training and validation. The evaluation scheme is presented in Figure 2B.

**Figure 2.**
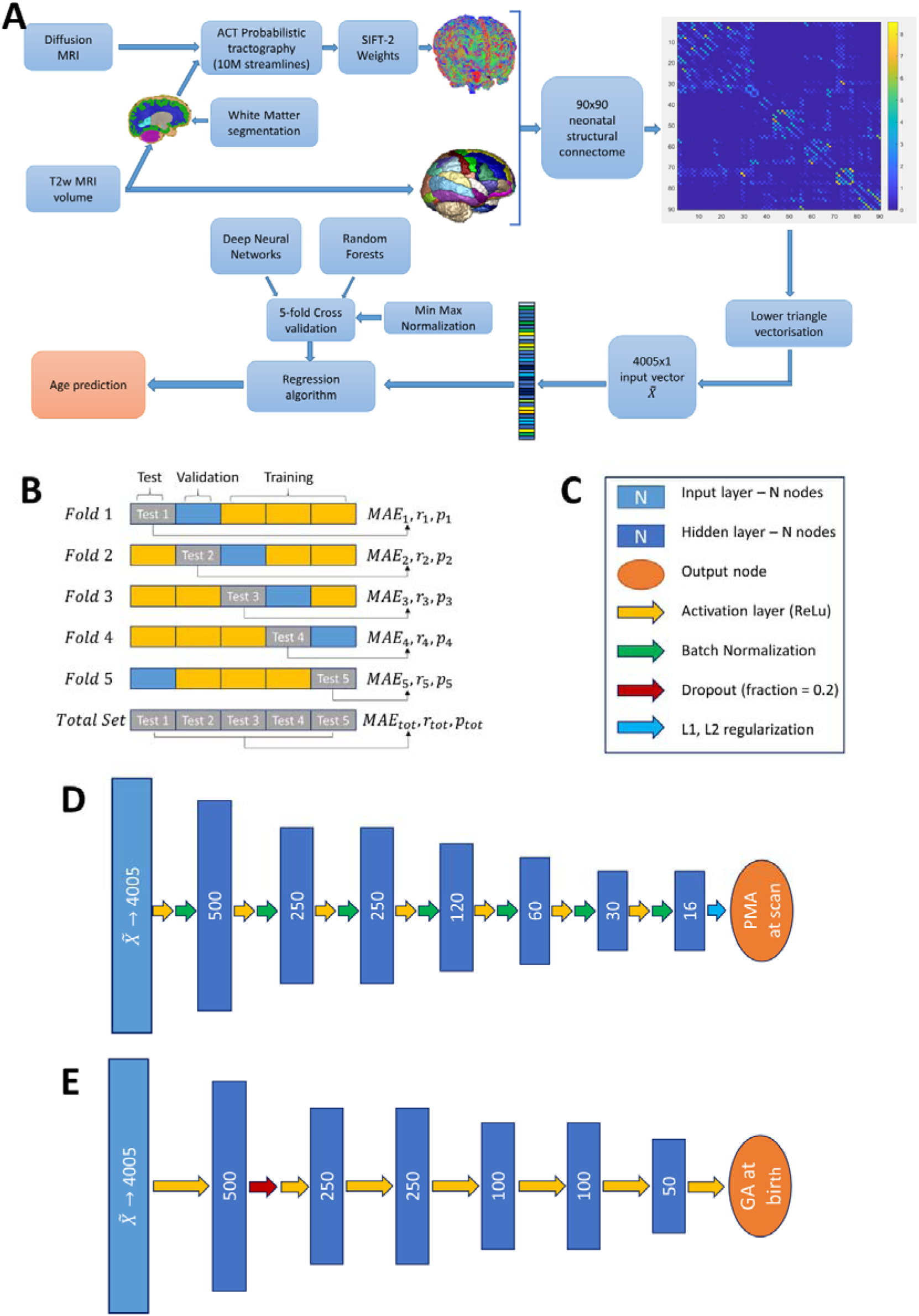
(A) Pipeline for age prediction from MRI. (B) Cross Validation protocol. (C) Legend for DNN architecture components. (D) PMA at scan DNN architecture. (E) GA at birth DNN architecture.

We split the data into k=5 groups (folds), with 20% of data used for testing at each fold. The remaining 80% of the data were further split for training (65%) to fit the models and validation (15%) to tune the hyperparameters. Min-max normalisation presented in section 2.4.1 is fitted on the training/validation set, where normalization parameters are saved and applied to the test set.

We added a bias correction as previously described (Smith et al. 2019; Peng et al. 2019) to correct age dependency of the training residuals. Briefly, we used a linear model *Y*′ *= f (X) = α Y + β* to obtain an unbiased estimate of *Y*′ as 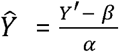, where the parameters *α* and *β* are estimated during training (on both the combination of training and validation set) and are thus applied directly to the test set. We obtained our final corrected prediction *Ŷ*_*i*_ for each structural connectome as follows:

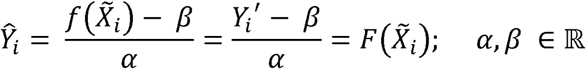

The final performance is calculated by averaging test-set performance over the 5 folds. We used mean absolute error (MAE) as our evaluation metric, calculated on each test set *k* as follows:

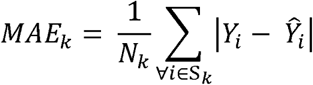

Where *N*_*k*_ is the number of subjects belonging to test set *k (*S_*k*_) and *Y*_*i*_ and *Ŷ*_*i*_ are actual and predicted outcome of subject *i*. In addition, we also evaluate MSE and *R*^2^ scores for each test set k, which are calculated as follows:

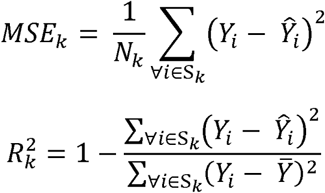

Where 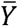 is the mean actual age of test set *k*. We also calculated Pearson’s Correlation (*r*_*k*_) and p-value (*p*_*k*_)between actual (*Y*) and predicted output (*Ŷ*) for each test set. As we obtain a prediction for every subject (albeit with different models) we can compute the *MAE*_*tot*_, *MSE*_*tot*_,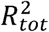, *r*_*tot*_ and *p*_*tot*_ by considering the predictions of the 5 test sets (see Figure 2A). Finally, we assessed the presence of heteroscedasticity in our predictions by comparing the variance 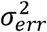 of the absolute error – a lower variance signifies more homoscedastic predictions.

#### 2.4.6 Prediction of PMA at scan in term-born infants

To build a model of typical development of connectivity we used the full cohort of 418 term-born babies (GA at birth >= 37) with PMA at scan between 37 and 45 weeks.

We first predicted PMA at scan from the vectorised and normalized structural connectome 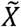 using RF regressor model. Optimal parameters of the model (max depth = 250, number of estimators = 30) were found by performing a grid search in an independent dataset (see section 2.4.3). We trained each fold on N≈335 samples (80%) including a validation set. We then tested the model on the remaining set (N≈83, 20%) in each fold, thus being able to predict age at scan on all 418 structural connectomes of term infants (see Figure 2A).

In a similar fashion, we also trained a regression DNN to predict PMA at scan from the vectorised and normalized structural connectome 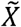. This DNN comprises one input layer with 4005 input nodes, 7 hidden layers, 6 activation layers (ReLu), 5 batch normalisation layers and one output layer with one node. Training was done for 50 epochs with learning rate of 0.007 and remaining parameters with default value. Detailed structure of the architecture of this DNN is provided in Figure 2D.

We applied the previously described bias correction method on both DNN and RF, by fitting *α* and *β* for each model *f*_*k*_ using both the training and validation set; thus reaching 5 distinct models *F*_1_,*F*_2_,…,*F*_5_ for both the DNN and RF methods.

#### 2.4.7 Prediction of GA at birth

To assess the effect of preterm birth on structural connectivity we trained a prediction model for GA at birth from 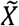 with both DNN and RF in a similar fashion as previously described for prediction of PMA at scan.

Since the dHCP cohort has significantly more term-born than preterm-born infants, there is a “class imbalance” in the GA distribution that may skew the model prediction. We therefore randomly selected a sub-sample of term subjects that had, on average, equal density of subjects on each GA at birth weekly bin. Our 106 preterm infants were distributed in 15 different GA at birth bins (22w-23w; 23w-24w … 36w-37w), thus providing an average of 7 infants per age category. We kept all 106 preterm infants and randomly sampled 7 infants from each of the term age categories (37w-38w, 38w-39w, … 41w-42w) and the 4 subjects born 42w-43w (as only 4 were available between 42w and 43w), for a total of 39 term infants. This resulted in a total of 145 infants with a balanced distribution (see Figure 1C).

We first attempted prediction of GA at birth from the vectorised and normalized structural connectome 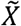 using RF with optimal parameters max depth = 300, number of estimators = 50. For each fold, we trained using N≈116 samples (80%) including a validation set. We then tested the model on the remaining set (N≈29, 20%) in each fold, predicting GA at birth for all 145 structural connectomes considered.

In a similar fashion, we also trained a DNN to predict GA at birth. This DNN consists of one input layer, 6 hidden layers, 6 activation layers (ReLu), one dropout layer, and one output layer. 120 epochs were used for training, with learning rate 0.003 and remaining parameters at default value. Detailed information on the architecture is provided in Figure 2E.

We applied the bias correction method on both DNN and RF by fitting *α* and *β* for each model *h*_*i*_ from the validation set; thus reaching 5 distinct models *H*_1_,*H*_2_,…*H*_5_ for both the DNN and RF methods.

#### 2.4.8 Brain maturation index

We defined brain maturation index *δ* (also called brain age or predicted age difference in the literature) as the difference between the predicted age *Ŷ* and true age *Y* of a subject n (Dosenbach et al., 2010): *δ*_*i*_ = *Ŷ*_*i*_ − *Y*_*i*_

We developed a model of typical brain development by training 5 models to predict PMA at scan on term-born infants only (section 2.4.6). Prediction of PMA at scan for each preterm subject was computed by taking the mean of the predictions from each of the 5 DNN trained models *F*_*k*_ from each cross-validation partition:

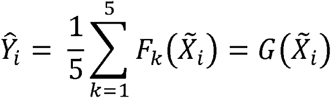

Following this, we computed the brain maturation index *δ*_*i*_ of each preterm subject:

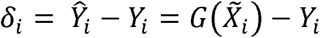

### 2.5 Sensitivity analysis to obtain feature relevance

The interpretation of models was computed using feature sensitivity analysis (Saltelli 2002) in a similar fashion for both the RF and DNN models on both predictions of GA and PMA (thus obtaining 4 different interpretations or “relevance maps”). We iteratively zeroed out every one of the 4005 connections and computed the magnitude of the change in predictions when zeroing out each feature compared to prediction from the unaltered connectome. The higher the magnitude of the change, the more impacted the model is to the given feature in the computing predictions. For each model, this was computed across all subjects and all folds; we took the mean magnitude of change across all folds as final feature of importance. We thereby obtained for each model features of importance for each of the 4005 connections. Following this step, we were also able to identify the most relevant brain regions, by summing the relevance across each column of each of the 4 relevance maps obtained. We used BrainNetViewer (Xia et al, 2013) to visualize the different relevance maps.

### 2.6 Statistical methods

Differences between term and preterm cohorts on all relevant characteristics were assessed by computing a two tailed independent t-test or chi-square test as appropriate. The association between brain maturation index *δ*_*i*_ and neurodevelopmental outcomes was assessed with Pearson’s Correlation coefficient for all preterm infants having both *δ*_*i*_ and BSID-III developmental outcomes at age 18 months corrected age. All outcomes were corrected for socio economic status, captured by the English Index of Multiple Deprivation (IMD) Rank. The IMD is a geographical measure that summarizes information from 38 different factors such as income, employment, education, crime rates and health situation for all postcode areas in England (Index of Multiple Deprivation, 2015). A lower IMD rank indicates a lower level of deprivation. We used the mother’s postcode at the time of birth to calculate this. All p-values presented are uncorrected for multiple comparisons

### 2.7 Data availability

The imaging and collateral data from the dHCP can be downloaded by registering at https://data.developingconnectome.org/

Structural connectivity networks and code used to predict age at birth and age at scan are available in https://github.com/CoDe-Neuro/Predicting-age-and-clinical-risk-from-the-neonatal-connectome

## 3. Results

### 3.1 Sample characteristics

There were no significant differences in PMA at scan and male/female proportion between term and preterm neonates in this study. For the subjects for which 18 months BSID-III follow-up neurodevelopmental assessment was available, there were no significant differences in outcomes between term and preterm infants. However, a significant group difference (p=0.002) was found in IMD scores, with term infants showing significantly higher deprivation than preterm infants. Detailed cohort characteristics are provided in Table 1 and Figure 1. Neurodevelopmental outcome details are provided in Table 1.

### 3.2 Prediction of PMA at scan

#### 3.2.1 Prediction of PMA at scan with RF

We trained a RF regressor to fit for PMA at scan from vectorised and normalized structural connectome 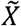 on term infants only. We obtained *MAE*_*tot*_ = 0.84 weeks,*MSE*_*tot*_ =1.10, 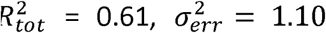 with correlation between true and predicted *r*_*tot*_ =0.79 *p*_*tot*_ < 0.001.Figure 3A shows true PMA vs predicted PMA on each of the 5 cross-validation folds. Detailed results of each fold are presented in Figure 3C.

**Figure 3.**
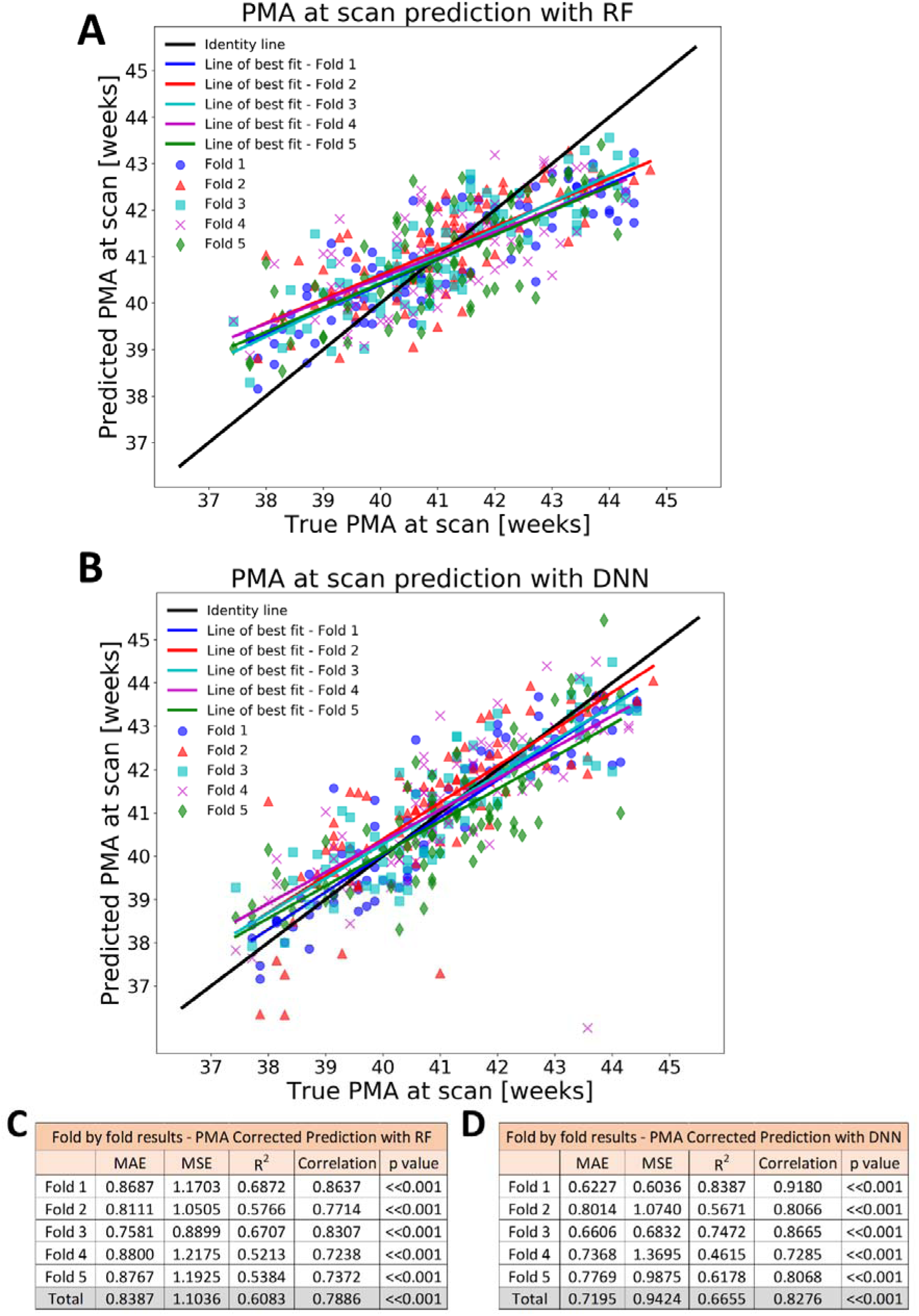
Detailed results of prediction of PMA at scan on term cohort. True vs Predicted with (A) RF, (B) DNN. Fold by fold result with (C) RF, (D) DNN.

#### 3.2.2 PMA at scan prediction with DNN

Similarly, we trained a DNN regressor to fit for PMA at scan from vectorised and normalized structural connectome 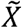 on term infants only. We obtained *MAE*_*tot*_ = 0.72 weeks, *MSE*_*tot*_ = 0.94, 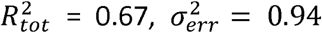 with correlation between true and predicted *r*_*tot*_ = 0.82 and *p*_*tot*_ <0.001. Figure 3B shows true PMA vs predicted PMA on each of the 5 cross-validation folds. Detailed results of each fold are presented in Figure 3D.

### 3.3 Prediction of GA at birth

#### 3.3.1 Prediction of GA at birth with RF

We trained a RF regressor to fit GA at birth from vectorised and normalized structural connectome 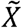 on balanced data (145 infants). We obtained *MAE*_*tot*_ = 2.76 weeks, *MSE*_*tot*_ = 12.95, 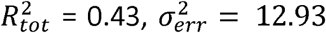 with correlation between true and predicted *r*_*tot*_ =0.67 and *p*_*tot*_ < 0.001. Figure 4A shows true GA at birth vs predicted GA at birth on each of the 5 folds fold. Detailed results of each fold are presented in Figure 4B.

**Figure 4.**
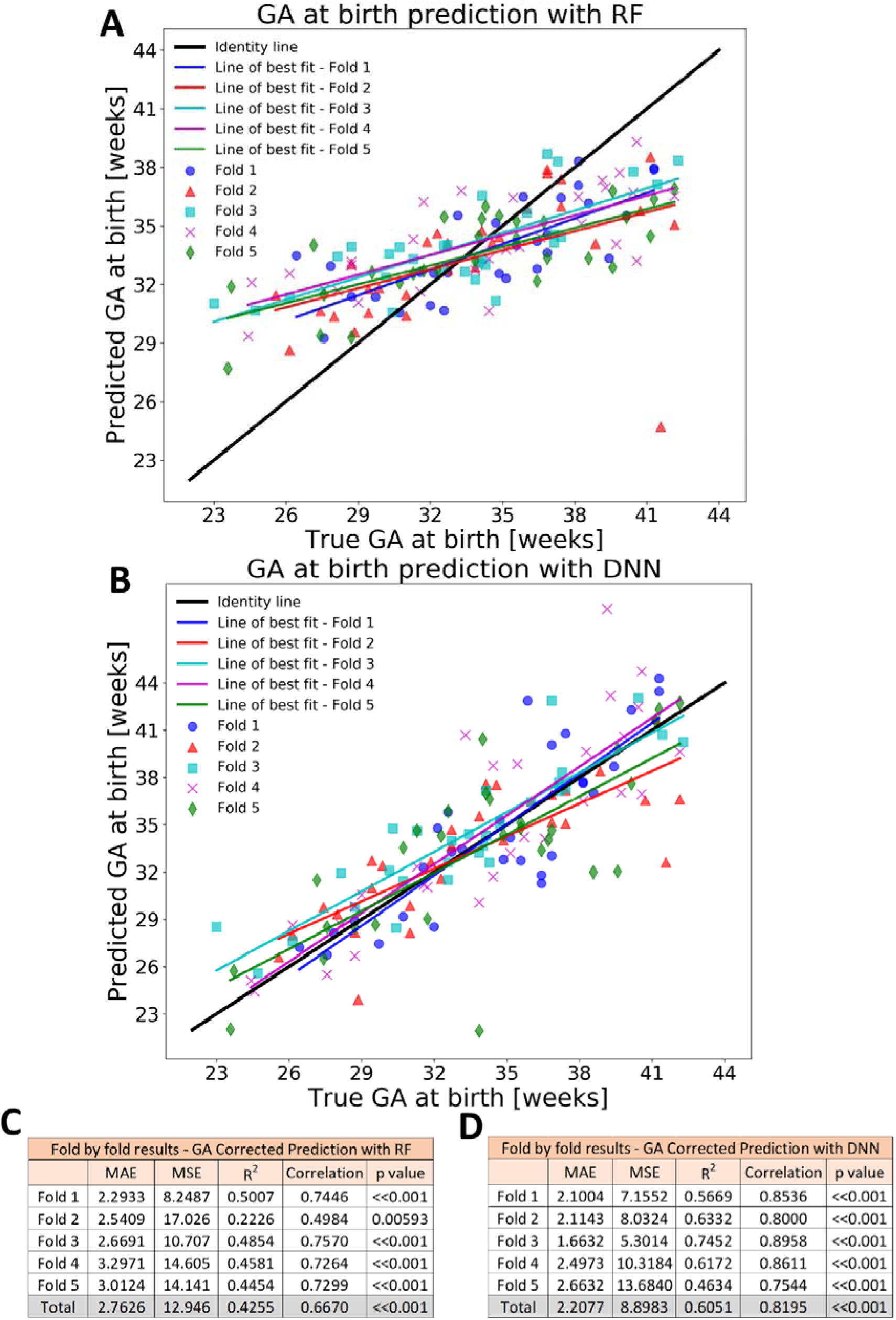
Detailed results of prediction of GA at birth. True vs Predicted with (A) RF, (B) DNN. Fold by fold result with (C)RF, (D)DNN.

#### 3.3.2 Prediction of GA at birth with DNN

Similarly, we trained a DNN from vectorised and normalized structural connectome 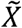 on balanced data. We obtained *MAE*_*tot*_ = 2.21 weeks, *MSE*_*tot*_ = 8.90, 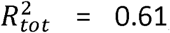, 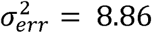 with correlation between true and predicted *r*_*tot*_ = 0.82 and *p*_*tot*_< 0.001 Figure 4B shows true GA vs predicted GA on each of the 5 folds. Detailed results of each fold are presented in Figure 4D.

### 3.3 Sensitivity analysis

We computed sensitivity analysis on each of the 4 models as described in the methods section, thereby obtaining 4 distinct connection relevance maps. From the relevance maps, we identified the 5 most relevant connections and brain regions (nodes) from each model, which are shown in Figure 6 C, D, G and H. Detailed relevance maps are shown in Figures 6 A, B, E and F). For visualization purposes, we computed and showed the z-score of each connection and region relevance instead of the raw relevance (magnitude of change) computed and show the 80 connections with higher relevance for each model (∼2 centile).

Visualisation of relevant connections shows different patterns across models, though similar types of connections and regions were detected as important. The thalamus, temporal lobe, frontal cortex, cingulate gyrus and putamen regions were identified as particularly important regions in all four models and were also involved in relevant connections. Connections within the frontal cortex, and between the frontal cortex and temporal and parietal lobe, cingulate and precuneus were highly relevant. Though not statistically significant, right hemisphere nodes were on average more relevant across all 4 models.

### 3.4 Brain Maturation Index

We computed the predicted PMA at scan of each preterm-born infant scanned at term-equivalent age by averaging out the 5 predictions from the DNN term trained models. We obtained MAE of 1.01 weeks on prediction of 106 preterm infants (*MSE*_*tot*_ = 2.24,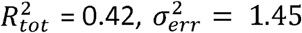), with correlation between true and predicted age between *r*= 0.79,(*p* < 0.001).True vs predicted age is presented in Figure 6A.

We computed the brain maturation index *δ*_*i*_ from each prediction. Brain maturation index (*δ*_*i*_) was significantly correlated with BSID-III gross motor scale at 18 months corrected age (*r* = 0.3751, *p* = 0.0008, Figure 6 B and C).

## 4. Discussion

We have used machine learning to uncover structural brain connectivity patterns associated with trajectories of early life development from high dimensionality neonatal MRI data. This enabled us to accurately predict PMA at scan in a large sample of term-born infants, and accurately predict GA at birth in a sample of preterm infants from scans at term equivalent age. The connectome features driving our predictions are of biological importance: in the preterm cohort we have shown significant correlation between a connectome derived brain maturation index and motor development at 18 months corrected age.

We achieved high accuracy in our prediction of PMA at scan in term born babies, reaching a low MAE of 0.72 weeks (*MSE*_*tot*_ = 0.94, 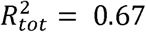) and a high correlation between true and predicted age (*r*_*tot*_ = 0.83; *p*_*tot*_ < 0.001) While the structural connectome presents several important challenges to study brain connectivity (Campbell and Pike, 2014) including high numbers of false positive streamlines (Maier-Hein et al., 2017), our results suggest that there is reliable information present to capture the subtle changes associated with weekly development. There have been several studies evaluating white matter microstructural and connectivity changes during the first days after birth. Indeed, it has been found that the postnatal period is marked by further dendritic arborization, refinement of existing intracortical connections, and an increase in synaptogenesis which results in an abundance of connections (for a review see (Keunen et al., 2017)). Some important changes have also been found in the structural connectome in the early postnatal period, mainly an increase in integration (the ease with which different brain regions communicate) and segregation (presence of clusters, i.e., capacity for specialised processing) (Batalle et al., 2017). These changes are likely captured by DNN and underlie the accurate prediction of PMA at scan. To our knowledge only a few studies have attempted the prediction of PMA at scan from the structural connectome. Using an RF method Brown and colleagues achieved prediction precision of 1.6 weeks using a sample of moderate and very preterm infants (77 scans, GA at birth between 24 and 32 weeks, scanned between 27 and 45 weeks) (Brown et al., 2017). Kawahara et al used a cohort of 115 preterm infants (born between 24 and 32 weeks gestational age) and reported a MAE of 2.17 weeks and r between true and predicted age of 0.87. Although widespread connections contributed to their predictive models, one (between the Right Lingual Gyrus and Fusiform Gyrus) was deemed to be of high importance (Kawahara et al., 2017). In addition to an improved MAE, using a sensitivity analysis approach we have shown a complex pattern of connections used for prediction of PMA (Figure 5A-D). The most important connections identified were thalamocortical or between frontal and temporal lobes, although widespread connections across the whole brain contributed to prediction. This is in keeping with a large body of previous literature on early life brain development: thalamocortical connections are developing late in the third trimester (Batalle et al., 2018), and are sensitive to disruption due to preterm birth or perinatal pathology (Ball et al., 2012, Ball et al. 2013, Batalle et al., 2017). Thalamic maturation continues in the first postnatal days (Kostovic and Jovanov-Milosevic, 2015), and it is therefore not surprising that inter-individual differences in thalamic connectivity appear to be important in the prediction of PMA at scan. We identified the superior frontal cortex and frontal orbital gyrus as two of the most important nodes in our PMA at scan prediction, which is in line with literature showing that connections to these regions are developing rapidly at term equivalent age (van den Heuvel et al., 2015; Kawahara et al., 2017). Right sided connections and nodes appear slightly more predictive of PMA than those on the left (although this difference was not significant). Structural asymmetry is well established by term equivalent age, although right sided connections are less efficient (Ratnarajah et al., 2013). One previous group has suggested that right sided connections are more important in predictions of neurodevelopment (Girault et al., 2019) and it is reasonable to hypothesize that inter-individual variation in the rate of development may also be important in prediction of PMA.

**Figure 5.**
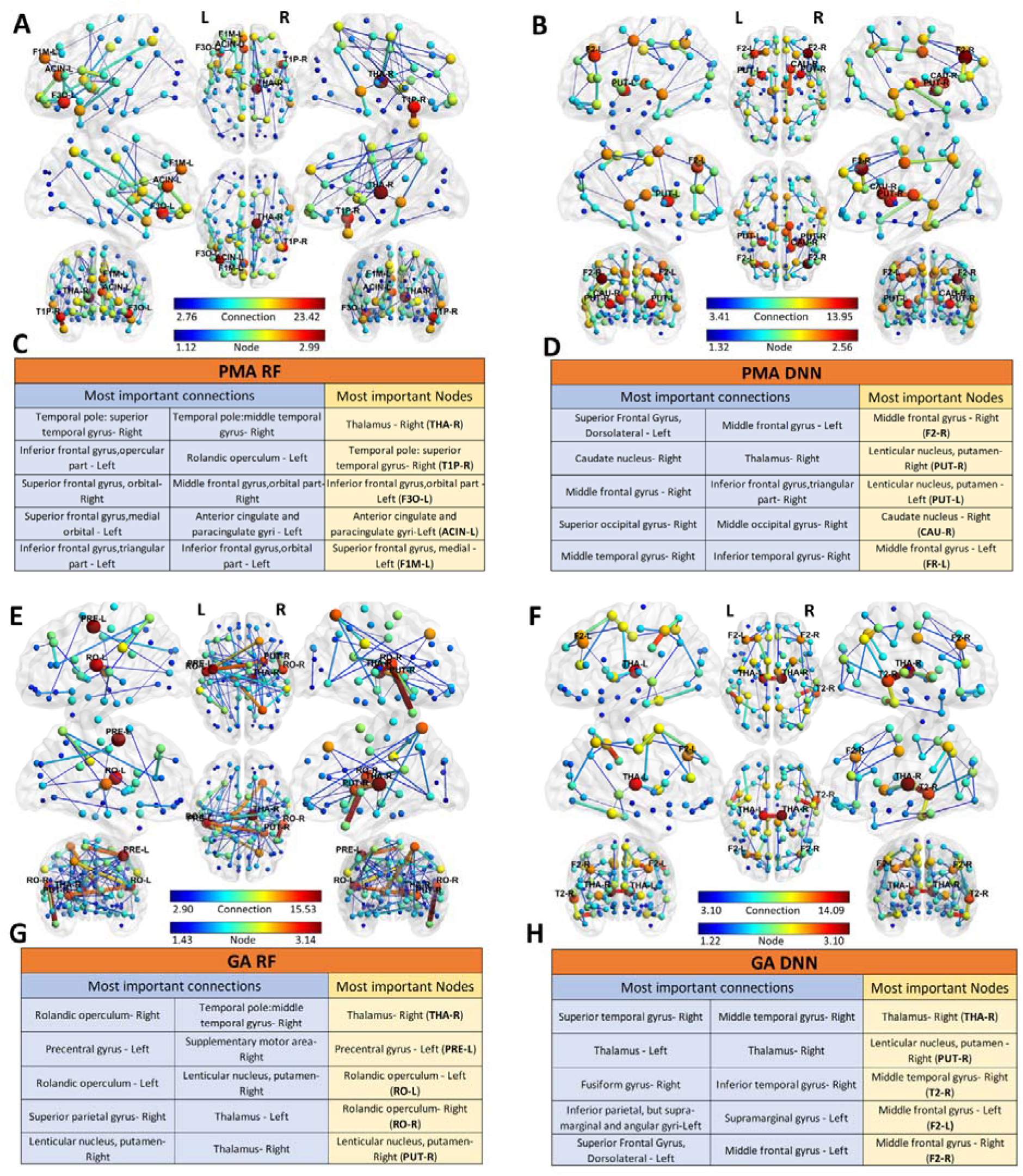
Relevance of connections and nodes for predictions of PMA with RF (A) and DNN (B), as well as GA with RF (E) and DNN (F); colour scale of each connection and region is the z-value of the computed relevance. Tables showing five most relevant connections and regions for predictions of PMA with RF (C) and DNN (D), as well as GA with RF (G) and DNN (H).

We have also demonstrated good ability to predict GA at birth from the neonatal connectome (DNN performance of *MAE*_*tot*_ = 2.21 *weeks*; *MSE*_*tot*_ *= 8*.*90;* 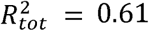 *r*_*tot*_ *= 0*.*82; p*_*tot*_ *< 0*.*001*). The nodes and connections of greatest importance to our models were mostly in and between the thalamus, frontal, and temporal lobes, and showed a slight right sided dominance (Figure 5E-H). The DNN model outperformed the RF model in prediction of GA at birth (Figures 4 A-D), which may be partially attributed to the difference in the fundamental mechanics of the models used and the high nonlinearity of the data. The effects of low GA at birth on structural brain development are widespread and complex: of particular relevance to our findings preterm birth has been associated with reduced structural (Ball et al., 2013) and functional thalamocortical connectivity (Toulmin et al., 2015). Organisation of fronto-temporal connectivity may also be impaired by preterm birth although the reported pattern of preterm associated changes is heterogeneous, including a higher proportion of cortical-cortical connections (Ball et al., 2014), changes in global network efficiency (Brown et al., 2014; Lee et al., 2019) and decreased fronto-limbic connectivity (de Almeida et al., 2021). It is worth noting that there is considerable overlap in the features we found to be more important for predicting PMA and GA at birth, and the critical regions used for prediction (thalamus, frontal and temporal lobes) are some of the most frequently cited in other studies of early life connectivity (van den Heuvel et al., 2015; Pandit et al., 2015). It may be the case that the rate of development (and thus potential for inter individual difference) is highest for these connections in the neonatal period, which may explain why they are most readily identified by machine learning techniques.

Brain maturation indices, which compare a predicted age to true age are a useful tool to characterise individual variations in the maturational trajectory of brain connectivity (Cao et al., 2015). In adults, a positive *δ* (predicted age > true age) is viewed as undesirable, and possibly indicative of the emergence of cognitive decline (Jonsson et al., 2019). In neonates 2015; however the opposite is likely true - a negative *δ* (predicted age < true age) may be indicative of developmental delay. We computed *δ* for all preterm infants (n=50) who had completed an 18-month follow-up visit and found a positive correlation between brain maturation index and BSID-III gross motor score (r=0.375, p<0.001). There is a large body of research on the effects of prematurity on motor development – lower GA at birth is universally associated with lower BSID-III motor scores (Greene et al., 2012; Velikos et al., 2015; Ahn et al., 2017). There is also some evidence that individual motor development differences in preterm children are in part due to individual brain connectivity profiles – diffusion measures in the cingulum (Schadl et al., 2017), corpus collosum (Lean et al., 2019), and global white matter diffusion tensor imaging measures (Girault et al., 2019) have all been associated with neurodevelopmental outcomes up to 2 years of age. Thalamocortical connectivity, key to the prediction of both PMA at scan and GA at birth in our study, has previously been associated with early life cognitive development (Ball et al., 2015; Eixarch et al., 2016). Two recent studies also using the dHCP dataset have used normative modelling, and specifically individual deviations from population growth trajectories, to associate imaging and outcome (Dimitrova et al., 2020; O’Muircheartaigh et al., 2020). The correlation observed here between our brain maturation index and cognitive development is therefore in keeping with existing knowledge and gives our model biological validity. However, since brain maturation index is obtained as the error in a model trained with term-born infants only, it is reasonable to expect larger model error (more negative *δ*) linked to reduced GA at birth (extreme prematurity), which is in turn associated with poorer neurodevelopmental outcomes. Hence, it is difficult to disentangle whether brain maturation index is just capturing features linked to the structural connectivity phenotype of preterm babies, or if it is capturing information related to motor outcome at 18 months that is embedded in the structural connectome at birth.

Our analysis uses well established machine learning techniques; however, it is important to be aware of their limitations. We have assessed the performance of RF and DNN for prediction of key developmental characteristics in a large sample of neonates. To our knowledge this is the first application of these techniques in a large cohort of neonates. A previous study predicting PMA from the structural connectome used a convolutional neural network approach (BrainNetCNN) to extract information from edge-to-edge, edge-to-node and node-to-node data from connectomes(Kawahara et al., 2017). The spatial distribution of adjacency matrices is not necessarily reflective of brain region locality and connectivity characteristics, so in this study, when predicting age directly from connectome data we instead chose to use RF and DNN, which don’t require data known to have local correlations. Using this approach, we achieved better performance compared to RF on age prediction from the neonatal structural connectome. Prediction of PMA at scan was highly accurate with RF (*MAE*_*tot*_ = 0.84; *MSE*_*tot*_ = 1.10; 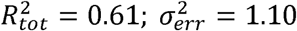; *r* = 0.79; *p* < 0.001), but DNN achieved better performance (*MAE*_*tot*_ = 0.72; *MSE*_*tot*_ = 0.94; 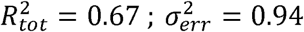; *r* = 0.83; *p* < 0.001), with a more homoscedastic distribution of prediction on each fold with DNN over RF was more of predictions over each of the 5 cross-validation folds (Figure 3). The improved performance of DNN over RF was more evident for prediction of GA at birth, with a better performance (*MAE*_*tot*_ *=2*.*21; MSE*_*tot*_ *=* 8.90; 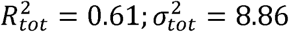; *r* = 0.82; *p* < 0.001) and more homoscedastic distribution of predictions on each fold with DNN over RF (*MAE*_*tot*_ *=2*.*76; MSE*_*tot*_ = 12.95; 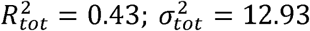; *r* = 0.67; *p* < 0.001) (Figure 4A-D). Another important consideration in machine learning MRI studies is prediction bias. As it can be observed in Figures 3A-B and 4A-B, prediction bias is still present after correction was applied (as described in section 2.4.5): since we fitted the bias correction values on the training set and blindly applied to the respective test sets (to prevent overfitting), the bias cannot be perfectly corrected. This is especially true for Random Forests – thus showing the higher stability and performance of DNN on unseen data. We chose to downsample the term cohort to achieve balance between age categories. Although this reduced our sample size, it allowed us to avoid a class imbalance problem, which could have caused a systematic positive bias for preterm infants (predicted GA > true GA). The relatively good performance of our model suggests that the impact of preterm birth on brain connectivity development is important and clearly apparent in the neonatal structural connectome. We achieved high performance in the prediction of PMA at scan in our preterm cohort by averaging the predictions from all 5 term trained DNN models (*MAE*_*tot*_ = 1.01; *MSE*_*tot*_ = 2.24 ; 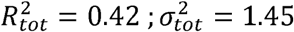; *r* = 0.79; *p* < 0.001). Although there was high accuracy in prediction and correlation between true and predicted age for preterm infants, predictions were on average inferior to those obtained for PMA at scan for term infants (Figure 6A). This is expected as preterm neonates are known to have specific differences in structural connectivity when compared with their term counterparts which may have reduced the generalizability of the predictive model (Ball et al., 2012; Batalle et al., 2017; Smyser et al., 2010). It is also important to mention that many methods have been proposed to interpret machine learning models and gain insight into the inner functioning of trained models (for a review, see (Molnar et al., 2020) and (Xie et al., 2020)). As we used both RF and DNN in this work, we chose to use a standard method of sensitivity analysis, as it could easily be applied in the same manner to both RF and DNN models instead of using other interpretation methods specifically tuned for each model. Furthermore, sensitivity analysis allows to visualize feature importance throughout the entire data set as opposed to in individual samples. This allowed us to easily compare network features contributing to the prediction of GA and PMA for both RF and DNN models.

**Figure 6.**
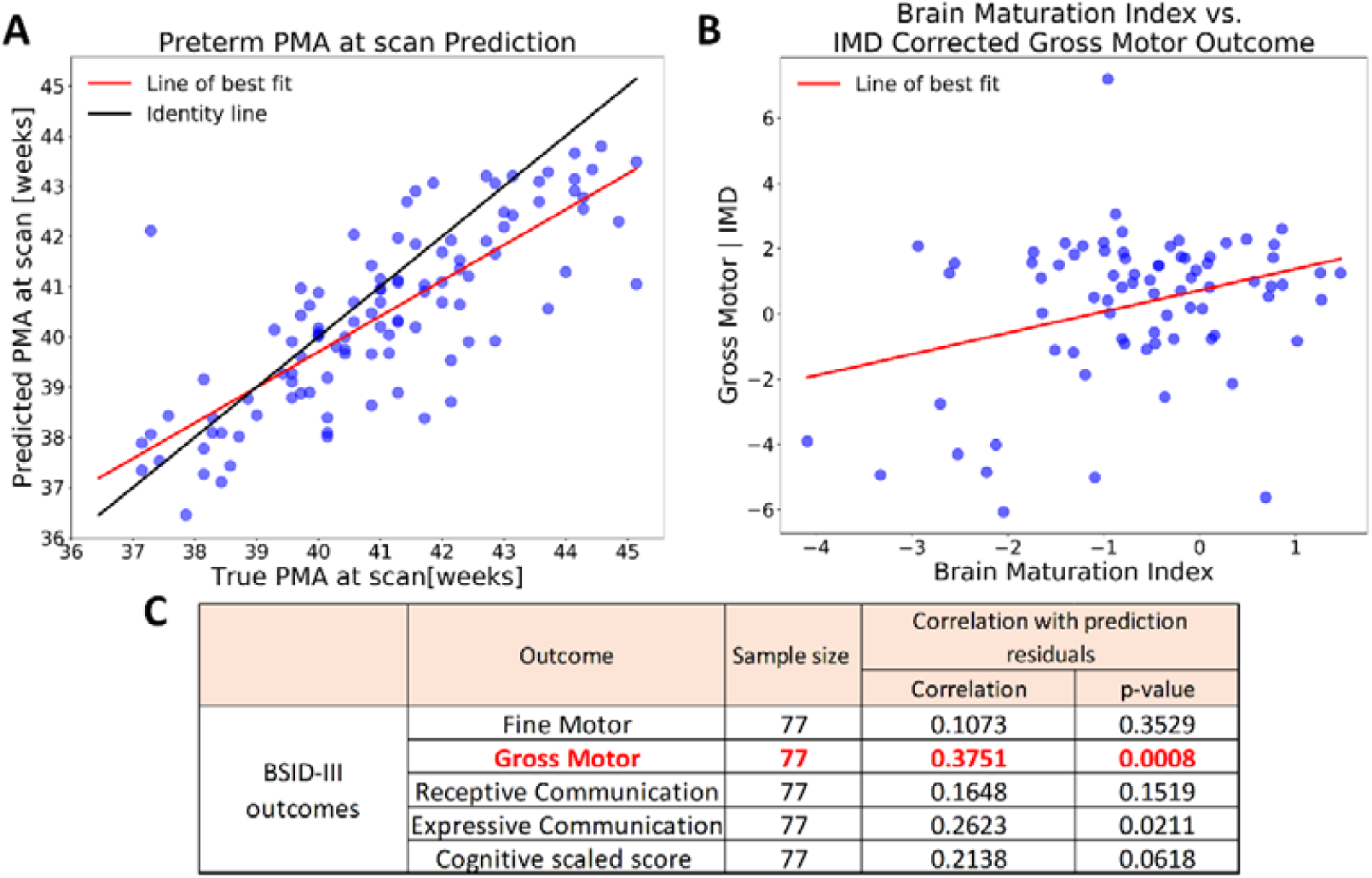
Association of Brain Maturation Index with BSID-III outcomes in preterm-born infants. (A) True vs predicted PMA at scan for preterm infants. (B) Brain maturation index δ vs BSID-III Gross motor outcome corrected for IMD. (C) Detailed correlation and p-values of preterm brain maturation index and BSID-III outcomes – statistically significant results surviving multiple comparisons correction are highlighted in bold red.

Although the dHCP dataset is one of the largest neonatal MRI resources currently available a key next step will be to investigate whether our results generalise to other cohorts. The regression algorithms were built and trained with the specific MRI acquisition protocol, brain parcellation and connectome generation methods developed for that project, which may limit how translatable our findings are. To enable this, our predictive algorithms have been made publicly available (https://github.com/CoDe-Neuro/Predicting-age-and-clinical-risk-from-the-neonatal-connectome). As the data was normalized prior to training, given a similar parcellation, we expect that significant correlation between true and predicted GA at birth or PMA at scan should be obtained if tested on different data. Future work could also focus on implementation of more modern machine learning approaches – for example geometric deep learning, which is specifically designed for data in non-Euclidean space (Bronstein et al., 2017). There is also increasing interest in predicting outcomes at age 18-24 months directly from neonatal brain connectivity, as done in (Girault et al., 2019). We have implemented our own version of their method and tested on the sub-set of our data set with available BSID-III at 18 months of age. However, no significant prediction capacity was reached with our data (data not shown). This might be due to different developmental outcome (Mullen scale instead of BSID-III), as well as differences in the pre-processing pipeline, or differences in the sample size and characteristics.

## 5. Conclusion

We have used DNN to accurately predict PMA and GA at Birth from the neonatal connectome and have used sensitivity analysis to describe the brain features most important in our models. We have additionally computed a brain maturation index which is associated with future motor development. We achieved a MAE of 0.72 weeks in predicting PMA at scan, demonstrating that the neonatal structural connectome contains key developmental information. Furthermore, prediction of GA at birth with MAE of 2.21 weeks shows that the patterns characteristic of prematurity are clearly present in the neonatal connectome, and can be uncovered with machine learning approaches. Several connectivity patterns identified as relevant for our models are in line with findings from existing neurodevelopmental studies. Lastly, brain maturation index was significantly correlated to BSID-III motor outcome at 18 months, suggesting the potential of this approach to develop biomarkers for prediction of atypical neurodevelopment.

