## Supplementary Table 1 for "Predicting age and clinical risk from the neonatal connectome"

**Supplementary Table 1.** Anatomical cortical regions of interest used as nodes, corresponding to the AAL atlas.

| **Anatomical ROI** | **Label** |  | **Anatomical ROI** | **Label** |
| --- | --- | --- | --- | --- |
| Precentral gyrus | PRE |  | Lingual gyrus | LING |
| Superior frontal gyrus, dorsolateral | F1 |  | Superior occipital gyrus | O1 |
| Superior frontal gyrus, orbital | F1O |  | Middle occipital gyrus | O2 |
| Middle frontal gyrus | F2 |  | Inferior occipital gyrus | O3 |
| Middle frontal gyrus,orbital part | F2O |  | Fusiform gyrus | FUSI |
| Inferior frontal gyrus,opercular part | F3OP |  | Postcentral gyrus | POST |
| Inferior frontal gyrus,triangular part | F3T |  | Superior parietal gyrus | P1 |
| Inferior frontal gyrus,orbital part | F3O |  | Inferior parietal, but supramarginal and angular gyri | P2 |
| Rolandic operculum | RO |  | Supramarginal gyrus | SMG |
| Supplementary motor area | SMA |  | Angular gyrus | AG |
| Olfactory cortex | OC |  | Precuneus | PQ |
| Superior frontal gyrus, medial | F1M |  | Paracentral lobule | PCL |
| Superior frontal gyrus,medial orbital | F1MO |  | Caudate nucleus | CAU |
| Gyrus rectus | GR |  | Lenticular nucleus, putamen | PUT |
| Insula | IN |  | Lenticular nucleus, pallidum | PAL |
| Anterior cingulate and paracingulate gyri | ACIN |  | Thalamus | THA |
| Median cingulate and paracingulate gyri | MCIN |  | Heschl gyrus | HES |
| Posterior cingulate gyrus | PCIN |  | Superior temporal gyrus | T1 |
| Hippocampus | HIP |  | Temporal pole: superior temporal gyrus | T1P |
| Parahippocampal gyrus | PHIP |  | Middle temporal gyrus | T2 |
| Amygdala | AMYG |  | Temporal pole:middle temporal gyrus | T2P |
| Calcarine fissure and surrounding cortex | V1 |  | Inferior temporal gyrus | T3 |
| Cuneus | Q |  |  |  |
